# Microbiota analysis of rural and urban surface waters and sediments in Bangladesh identifies human waste as driver of antibiotic resistance

**DOI:** 10.1101/2021.02.04.429629

**Authors:** Ross Stuart McInnes, Md. Hassan uz-Zaman, Imam Taskin Alam, Siu Fung Stanley Ho, Robert A. Moran, John D. Clemens, Md. Sirajul Islam, Willem van Schaik

## Abstract

In many low- and middle-income countries antibiotic resistant bacteria spread in the environment due to inadequate treatment of wastewater and the poorly regulated use of antibiotics in agri- and aquaculture. Here we characterised the abundance and diversity of antibiotic-resistant bacteria and antibiotic resistance genes in surface waters and sediments in Bangladesh through quantitative culture of Extended-Spectrum Beta-Lactamase (ESBL)-producing coliforms and shotgun metagenomics. Samples were collected from highly urbanised settings (*n* = 7), from rural ponds with a history of aquaculture-related antibiotic use (*n* = 11) and from rural ponds with no history of antibiotic use (*n* = 6). ESBL-producing coliforms were found to be more prevalent in urban samples than in rural samples. Shotgun sequencing showed that sediment samples were dominated by the phylum Proteobacteria (on average 73.8% of assigned reads), while in the water samples Cyanobacteria (on average 60.9% of assigned reads) were the predominant phylum. Antibiotic resistance genes were detected in all samples, but their abundance varied 1,525-fold between sites, with the highest levels of antibiotic resistance genes being present in urban surface water samples. We identified an IncQ1 sulphonamide resistance plasmid ancestral to the widely studied RSF1010 in one of the urban water samples. The abundance of antibiotic resistance genes was significantly correlated (R^2^ = 0.73; *P* = 8.9 × 10^−15^) with the abundance of bacteria originating from the human gut, which suggests that the release of untreated sewage is a driver for the spread of environmental antibiotic resistance genes in Bangladesh, particularly in highly urbanised settings.

**Importance:** Low- and middle-income countries (LMICs) have higher burdens of multidrug-resistant infections than high-income countries and there is thus an urgent need to elucidate the drivers of the spread of antibiotic-resistant bacteria in LMICs. Here we study the diversity and abundance of antibiotic resistance genes in surface water and sediments from rural and urban settings in Bangladesh. We found that urban surface waters are particularly rich in antibiotic resistance genes, with a higher number of them associated with plasmids indicating that they are more likely to spread horizontally. The abundance of antibiotic resistance genes was strongly correlated with the abundance of bacteria that originate from the human gut, suggesting that uncontrolled release of human waste is a major driver for the spread of antibiotic resistance in the urban environment. Improvements in sanitation in LMICs may thus be a key intervention to reduce the dissemination of antibiotic resistant bacteria.

## Introduction

The prevalence of antibiotic-resistant bacteria causing infections is increasing globally, but the clinical issues, including significant morbidity and mortality, posed by these bacteria are particularly alarming in low- and middle-income countries (LMICs) (1–4). Proposed drivers for the high burden of drug-resistant infections in LMICs include the unregulated sales of antibiotics and their misuse in clinical medicine, agriculture and aquaculture, an inadequate sewerage infrastructure, poor governance and low investments in health care (5, 6).

One of the challenges of studying AMR is to disentangle the spread of resistant bacteria and antibiotic resistance genes between humans, animals and the wider environment (7). For this reason, AMR is increasingly being studied from a collaborative and cross-disciplinary perspective that has been termed ‘One Health’ (8). The One Health concept for studying the spread of AMR is particularly relevant for LMICs due to the crucially important role of agriculture and aquaculture in the livelihoods of billions of people in many of these countries, especially the poorest ones (9). Asia is home to an estimated 74% of the world’s 570 million farms (10) and, in 2016, 89% of the global aquaculture production was estimated to originate from this continent (11). However, there are still major knowledge gaps on the spread of AMR in Asia from a One Health perspective.

Bangladesh is an LMIC in South Asia, where antibiotic-resistant infections are common among both hospitalised patients and the non-hospitalised population (12). The country has a number of unique characteristics that may contribute to the rapid spread of AMR. The capital city of Bangladesh, Dhaka, has a population of around 16 million people, with a population density that ranks among the highest of any megacity. Less than 20% of the households in Dhaka are connected to sewerage infrastructure (13), facilitating the spread of antibiotic-resistant bacteria via the environment. While a prescription is legally required to purchase antibiotics in Bangladesh, they can be readily acquired from many of the 200,000 drug stores across Bangladesh (14). In rural Bangladesh, aquaculture is widespread with more than 2 million tonnes of freshwater fish produced in 2017 from inland freshwater fisheries (15). A recent survey revealed that antibiotics are widely used in Bangladeshi aquaculture for disease prevention and growth promotion. The most prominent classes of antibiotics employed are the tetracyclines, but other antibiotic classes including β-lactams and sulphonamides are also used (16). The use of antibiotics in Bangladesh is regulated in line with the European Union standards for antibiotic use in aquaculture, but Bangladesh has been found to be in breach of these regulations several times (17). The causes of antibiotics overuse in aquaculture are multifactorial: pharmaceutical companies provide food which is premixed with antibiotics without the farmers’ knowledge, farmers administer antibiotics too often because they do not understand the instructions, and prophylactic use of antibiotics may be used to reduce the chance of damaging losses in production caused by disease (18). The combination of a densely populated country, intensive antibiotic usage in aquaculture and the potential for the dissemination of antibiotic-resistant bacteria through surface water thus provides a unique opportunity to study the spread of AMR from a One Health perspective.

In this manuscript, we use a combination of quantitative bacterial culture and metagenomic shotgun sequencing methods to disentangle pathways that contribute to the dissemination of antibiotic resistance. Specifically, we describe the abundance and diversity of microorganisms and antibiotic resistance genes in surface water in rural and urban settings in Bangladesh.

## Results

### Sample collection across urban and rural sites in Bangladesh

Freshwater surface water and sediment samples were collected from 24 sites across three districts in Bangladesh (Mymensingh, Shariatpur and Dhaka; Figure 1). These sites spanned both rural and urban areas with different population densities. Among rural sites, ponds used for aquaculture with a history of antibiotic use (*n* = 11) and ponds with no history of antibiotic use (*n* = 6) were sampled. Further information on sampling locations and protocols is provided in the Materials and Methods section and in Table S1. We used culture-dependent and culture-independent methods to study the abundance of antibiotic resistance genes and the diversity of microbiotas across the different sites.

**Figure 1.**
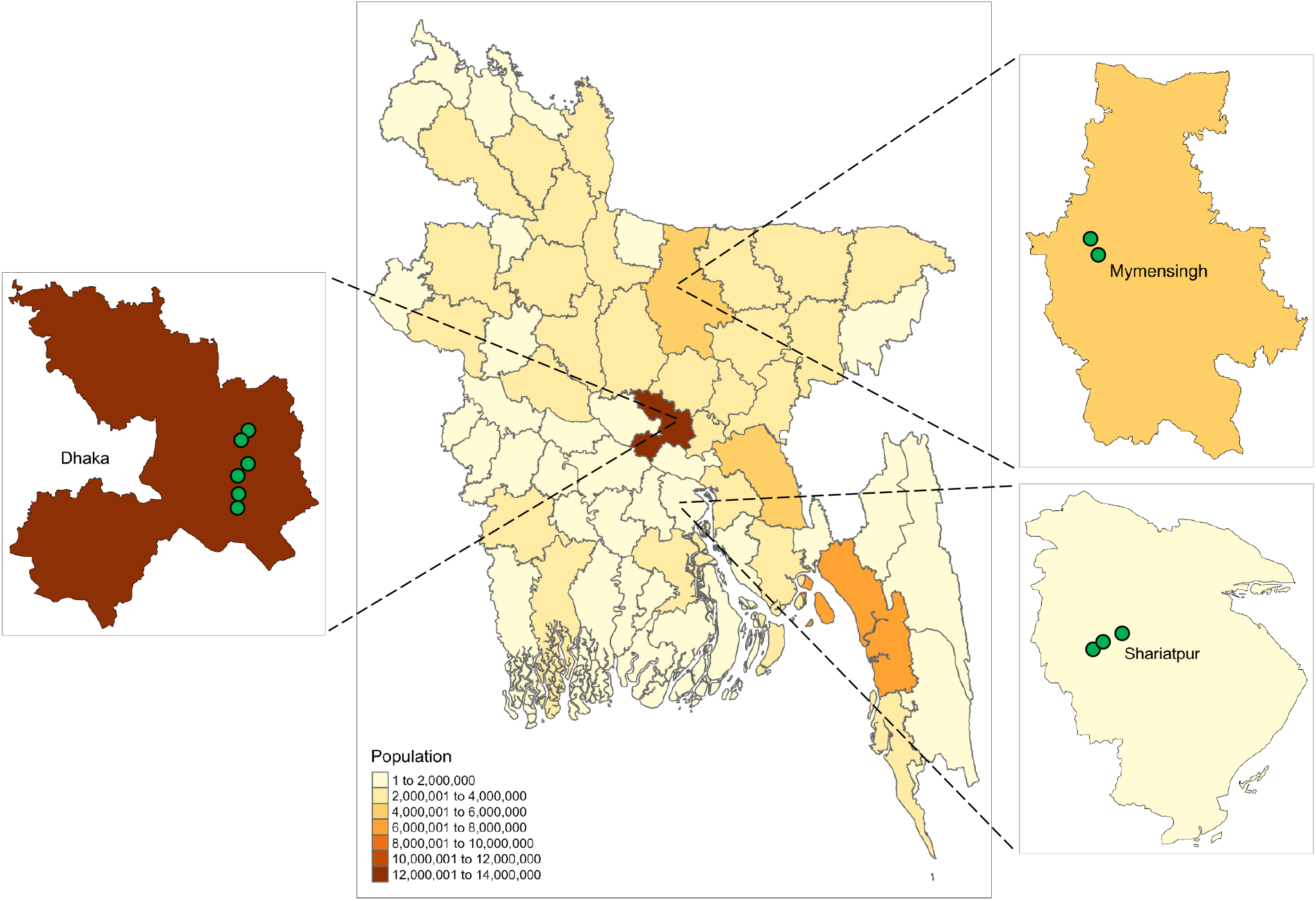
Map of Bangladesh showing the districts that the samples were collected from and the population of each district (obtained through https://data.humdata.org/dataset/bangladesh-administrative-level-0-3-population-statistics). Green circles represent sampling locations.

### ESBL-producing coliforms were more prevalent in urban samples than in rural samples

We quantitatively determined the burden of Extended Spectrum Beta-lactamase (ESBL) producing coliforms in the water and sediment samples from the different sampling locations and found that ESBL-producing coliforms were detected in significantly more urban samples (12/14) than rural samples (15/34) (Fisher exact test; *P* = 0.01). However, in samples that contained detectable levels of ESBL-producing coliforms there was no statistically significant difference in the viable counts of urban or rural samples (Figure 2).

**Figure 2.**
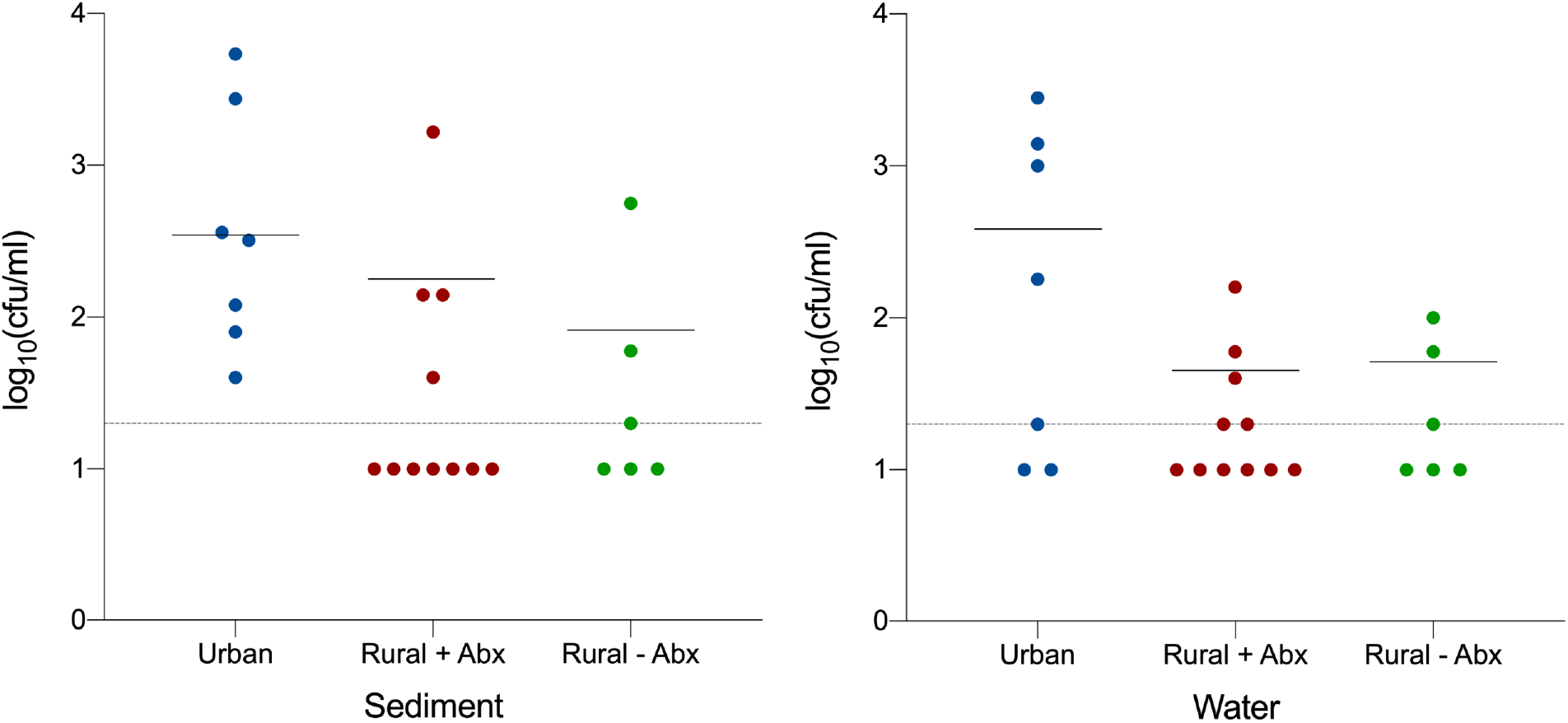
The abundance of ESBL producing coliforms, in log10 (cfu/ml), isolated from sediment and surface water in urban sites and rural settings with antibiotic use (+Abx) and without antibiotic use (-Abx) across Bangladesh. The horizontal dashed line represents the detection limit of 20 cfu/ml. Samples with ESBL-producing coliforms below the detection limit were plotted at log10 (cfu/ml) = 1.

### Microbiotas of surface water and sediments are distinct with higher levels of human gut bacteria in urban samples

Shotgun metagenomic sequencing was used to study the diversity and composition of the microbial communities in the different samples. An important determinant shaping the communities was the sample type, with distinct (PERMANOVA; *P* < 0.001) clustering of sediment and water samples (Figure 3A). The sediment samples were dominated by the phylum Proteobacteria (73.8%; standard deviation (SD) 27.1) while in the water samples Cyanobacteria (60.9%; SD 29.6) was the dominant phylum (Figure 3B). However, considerable variation in the composition of the microbial communities was observed as in five of the nine sediment samples collected in Mymensingh, the abundance of Euryarchaeota was greater than 50%, while in five Dhaka water samples Proteobacteria were present at levels greater than 45%. Water sample WAM6 had very high levels (>60%) of bacteriophage DNA. The sediment samples were dominated by typical soil bacteria such as *Pseudomonas*, *Azoarcus* and *Anaeromyxobacter* while the water samples were dominated by cyanobacteria such as *Cyanobium*, *Microcystis* and other typical aquatic bacterial species from the phyla Proteobacteria and Actinobacteria. Three bacteriophages (*Mycobacterium* phage rizal, *Microcystis aeruginosa* phage Ma LMM01 and an Epsilon15-like virus) were also identified at different sampling sites. It was apparent that many of the Dhaka water samples contained bacteria which are typically found within the gastrointestinal tract, including *Escherichia coli, Streptococcus infantarius, Bifidobacterium adolescentis* and *Prevotella copri*. Through microbial source tracking analysis of our shotgun sequencing data using the FEAST(19), we found that the urban water samples had a significantly greater (Kruskal-Wallis; *P* < 0.01) contribution from gut bacteria compared to the rural samples without previous antibiotic use (Figure 3C).

**Figure 3.**
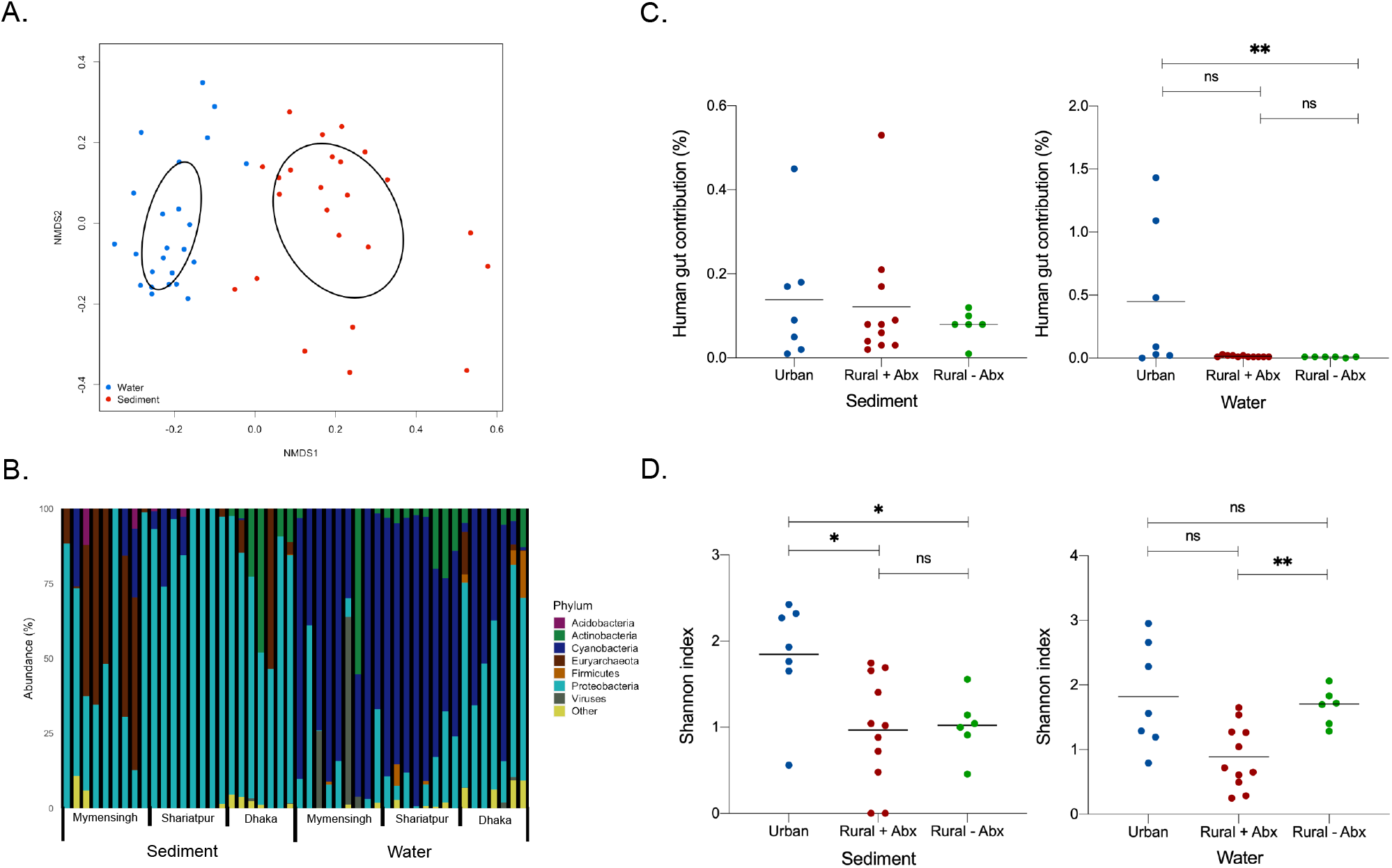
**A**. Non-metric multidimensional scaling (NMDS) analysis of a Bray Curtis distance matrix of species abundance. Stress 0.15. Ellipses represent standard deviation. **B.** Relative abundance (%) of Phyla across the 48 samples from sediment and surface water. **C.** Source-sink analysis, percentage contribution of human gut bacteria to the bacterial composition of the water and sediments samples. Kruskal-Wallis. **\*\****P* < 0.01. **D.** Shannon diversity values of species present in sediment and water samples from across Bangladesh. Brown-Forsythe ANOVA. **\****P* < 0.05 **\*\****P* < 0.005.

The urban sediment samples were significantly more diverse than both the rural samples with and without previous antibiotic use (Browne-Forsythe and Welch; *P* < 0.05). There was no significant difference in diversity between either of the rural sediment sample types (Figure 3D). On the other hand, the rural water samples without previous antibiotic use were significantly more diverse than the rural samples with previous antibiotic use (Browne-Forsythe and Welch; *P* < 0.005) but there was no significant difference between the urban water samples and either of the rural sample types.

### Urban samples carry the highest antibiotic resistance gene loads

A total of 114 different antibiotic resistance genes (ARGs) that confer resistance to 16 antibiotic classes were identified in the 48 samples from sediment and surface water. The urban samples had the greatest number of ARGs (n = 99) followed by the rural samples with previous antibiotic use (n = 49), while the rural samples with no previous antibiotic use had the fewest resistance genes (n = 36) (Figure 4). There was a large overlap between the ARGs present in the different sample types with the urban and rural + antibiotic samples sharing the greatest number of resistance genes (n = 24). There were 17 ARGs shared between all three samples types including five different beta-lactamase genes belonging to the *bla*_OXA_ and *bla*_RSA_ families.

**Figure 4.**
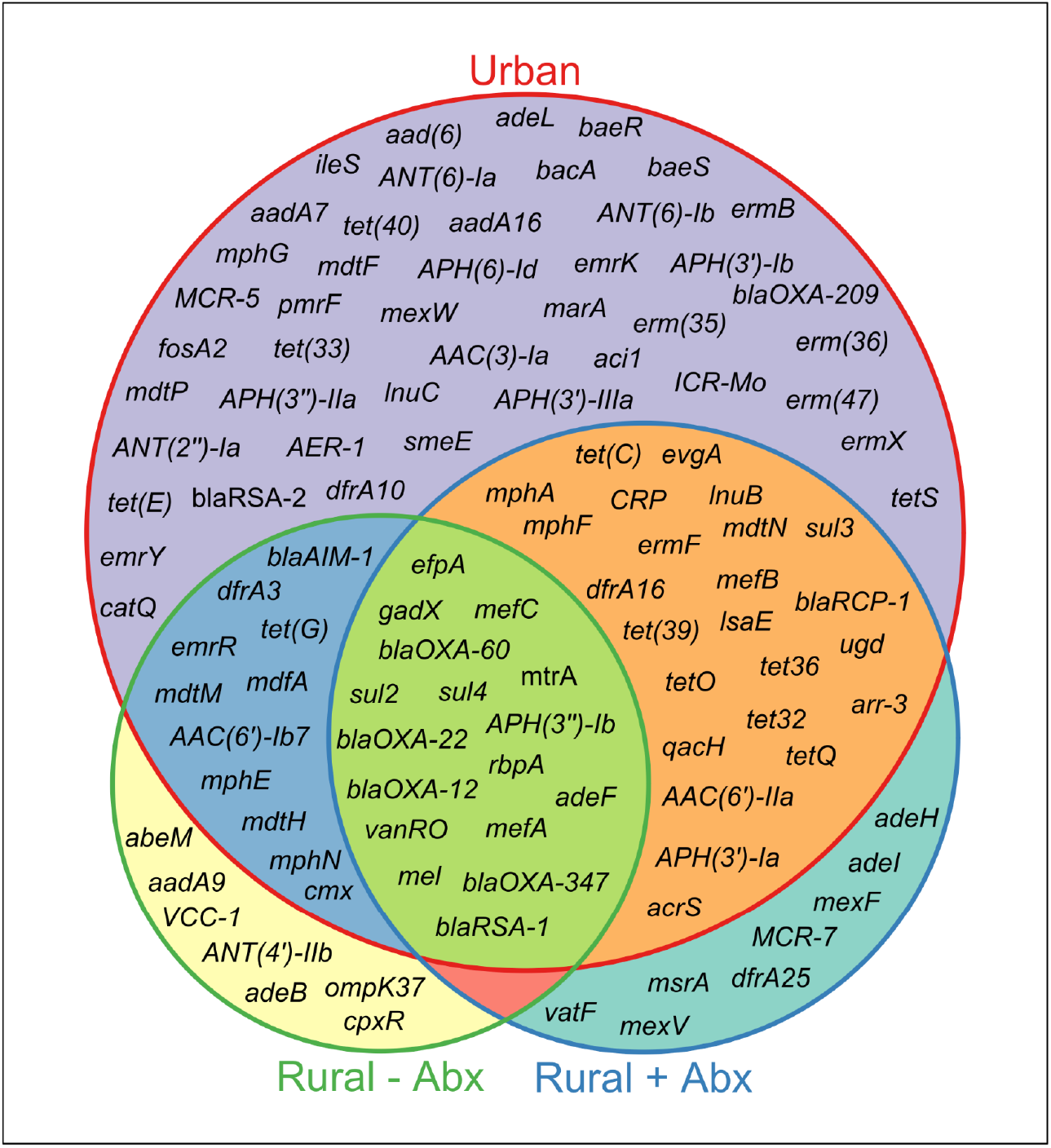
Distribution of antibiotic resistance genes across urban, rural without prior antibiotic use and rural with prior antibiotic use sample types. Circles are proportional to the number of antibiotic resistance genes present within each sample type.

The abundance of antibiotic resistance genes varied 1,525-fold between sites, with sample SAM6 (rural sediment sample with previous antibiotic exposure collected in Shariatpur) having the lowest abundance (0.078 Reads per Kilobase per Million reads [RPKM]) and sample WD7 (surface water sample collected in Dhaka) having the highest ARG abundance (120.45 RPKM). Of the paired sediment and water samples, the ARG abundance was on average 3 times greater in the water samples than the sediment samples (Wilcoxon; *P* < 0.0001). The urban sediment samples collected from around the city of Dhaka were found to have a significantly (Kruskal-Wallis; *P* < 0.05) greater total ARG abundance (median RPKM = 4.01, interquartile range (IQR) = 0.95 – 12.79) than the rural samples with prior antibiotic use (median RPKM = 0.60, IQR = 0.20 – 1.27) (Figure 5A). However, the urban sediment samples were not significantly different to the rural samples without antibiotic use (median RPKM = 0.72, IQR = 0.64 – 1.36). There was also no statistically significant difference (Kruskal-Wallis; *P* > 0.99) between ARG abundance in rural sediment with prior antibiotic use versus sediment from rural sites in which antibiotics had not been used. ARG levels in the water samples reflected that of the sediment samples, with the total ARG abundance in urban samples (median RPKM = 37.08, IQR = 5.71 – 97.74) being significantly higher (Kruskal-Wallis; *P* < 0.05) than the rural samples with previous antibiotic use (median RPKM = 4.30, IQR = 2.39 – 7.60) but not significantly different to the rural samples with no previous antibiotic use (median RPKM = 5.09, IQR = 1.80 – 11.68). As with the sediment samples there was no significant difference found between either of the rural sample types (Kruskal-Wallis, *P* > 0.99).

**Figure 5.**
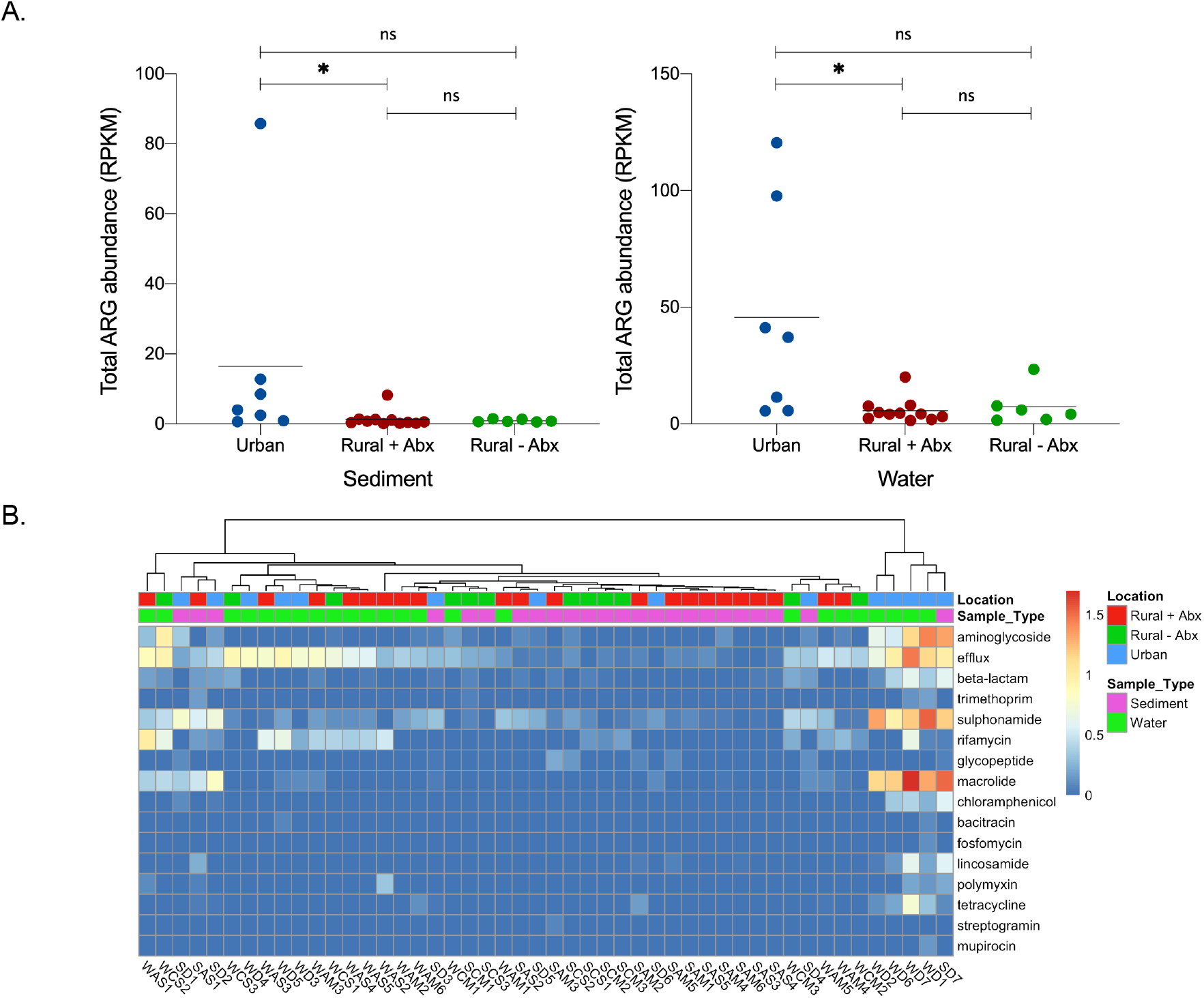
**A.** Abundance in RPKM of Antibiotic Resistance Genes (ARGs) in each sample (sediments and surface water; urban, rural with antibiotic use and rural without antibiotic use). Kruskal-Wallis test ^*^ *P* < 0.05. **B.** Heatmap representing the summed abundance (log10 transformed RPKM) of antibiotic resistance gene classes present in water and sediment samples from surface water sites across Bangladesh.

The individual antibiotic resistance genes were collated into 16 classes that cover resistance to specific antibiotics and a separate class for genes conferring antibiotic efflux mechanisms (Figure 5B). Efflux genes were present in 47 of 48 samples making it the most widespread ARG class. Other abundant antibiotic resistance classes were resistance to sulphonamides, macrolides and aminoglycosides. Urban water samples WD2, WD6, WD7 and WD1 and an urban sediment sample SD7 clustered together, with high levels of resistance genes from these classes.

### Abundance of human gut bacteria predicts levels of antibiotic resistance genes

There was a statistically significant correlation (R^2^ = 0.73; P = 8.9 × 10^−15^) between the aggregated abundance of ARGs and the levels of human gut bacteria across our study (Figure 6A). We also determined whether the levels of ESBL-producing coliforms are correlated with the total abundance of ARGs and observed a relatively weak but statistically significant correlation (R^2^ = 0.38; P = 1.8 × 10^−6^) (Figure 6B).

**Figure 6.**
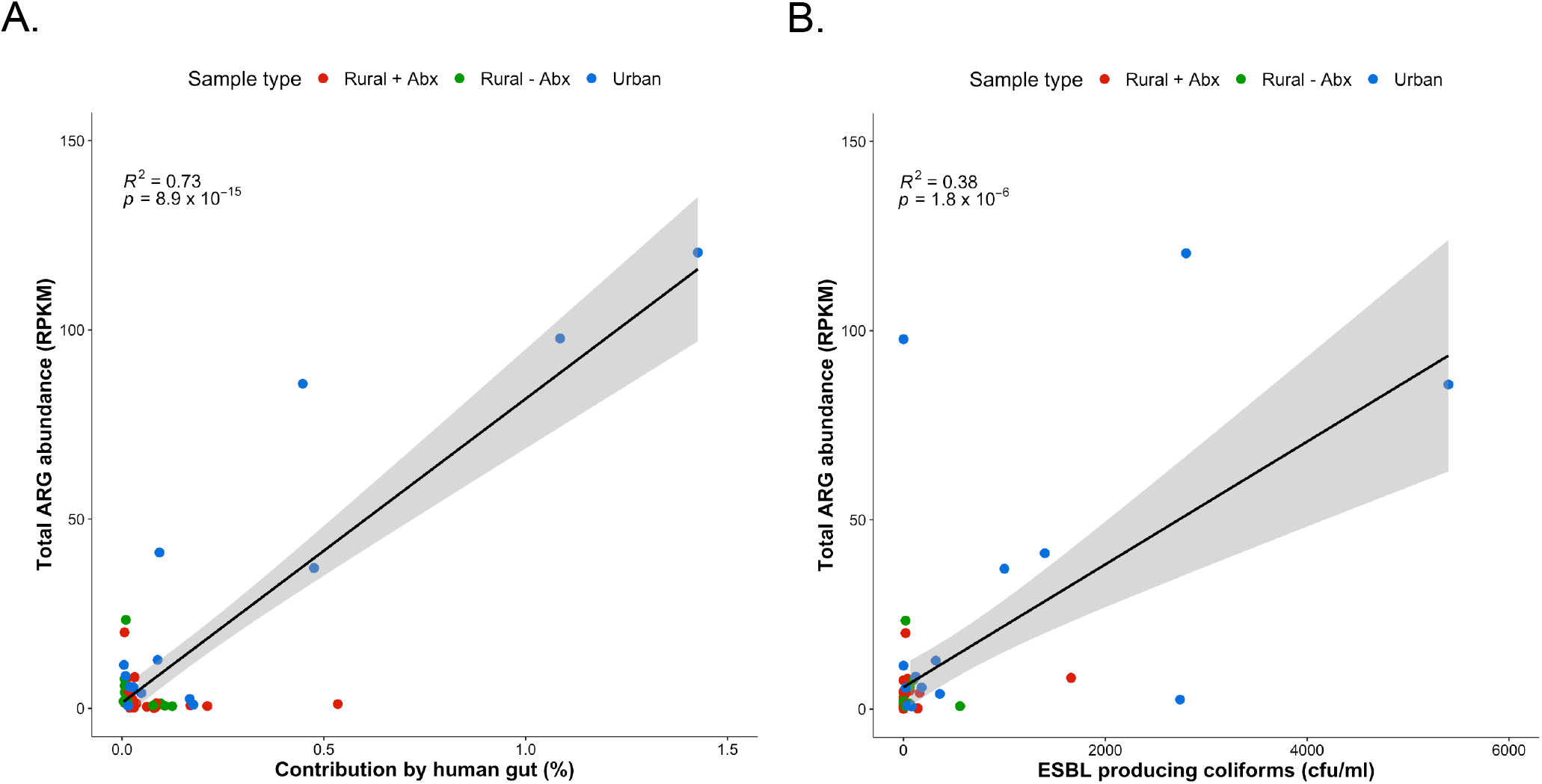
**A.** Correlation between the total antibiotic resistance gene (ARG) abundance (RPKM) and the percentage of bacteria contributed from the human gut within each sample. R^2^ = 0.73. *P* = 8.9 × 10^−15^. **B.** Correlation between the total ARG abundance (RPKM) and the number of ESBL producing coliforms (cfu/ml) in each sample. R^2^ = 0.38. *P* = 1.8 × 10^−6^. The grey area represents the 95% confidence interval.

### Urban sites were enriched in plasmids carrying antibiotic resistance genes

As antibiotic resistance genes were particularly abundant in water samples, we performed a metagenomic assembly of the short-read data from the surface water samples to recover complete plasmid sequences and study their potential association with antibiotic resistance. The metagenomic assemblies were queried against the PlasmidFinder database (20) to identify contigs which contained plasmid replication (*rep*) genes. Eleven contigs in our dataset contained *rep* genes (Table S2). Seven Gram-negative replicons were found which were related to representatives of the P and Q incompatibility groups or to small theta- or rolling circle-replicating plasmids. A single Gram-positive replicon, repUS43, was identified in sample WD1. Two plasmid contigs, k141_206349 (2113 bp) and k141_304072 (8535 bp), could be circularised (Figure S1). The latter plasmid, which we named pWD1, contained the sulphonamide resistance gene *sul2* adjacent to a complete copy of the mobile element CR2 (blue box in Figure 7), an IncQ1 replicon, three mobilisation genes (*mobABC*) and an origin-of-transfer (*oriT*). pWD1 was found to have 99.97% identity over 81% of its sequence to the canonical broad-host range mobilisable plasmid RSF1010 (21). Alignment and annotation of these two plasmids revealed that they were identical apart from in the region immediately downstream of *sul2*. In RSF1010 the insertion of the streptomycin resistance genes *strA-strB,* is associated with truncation of CR2 and the *rcr2* gene (Figure 7). While the RSF1010 configuration is common, the *sul2*-CR2 configuration in pWD1 was not found in any other IncQ1 plasmids in GenBank (searched December 9, 2020).

**Figure 7.**
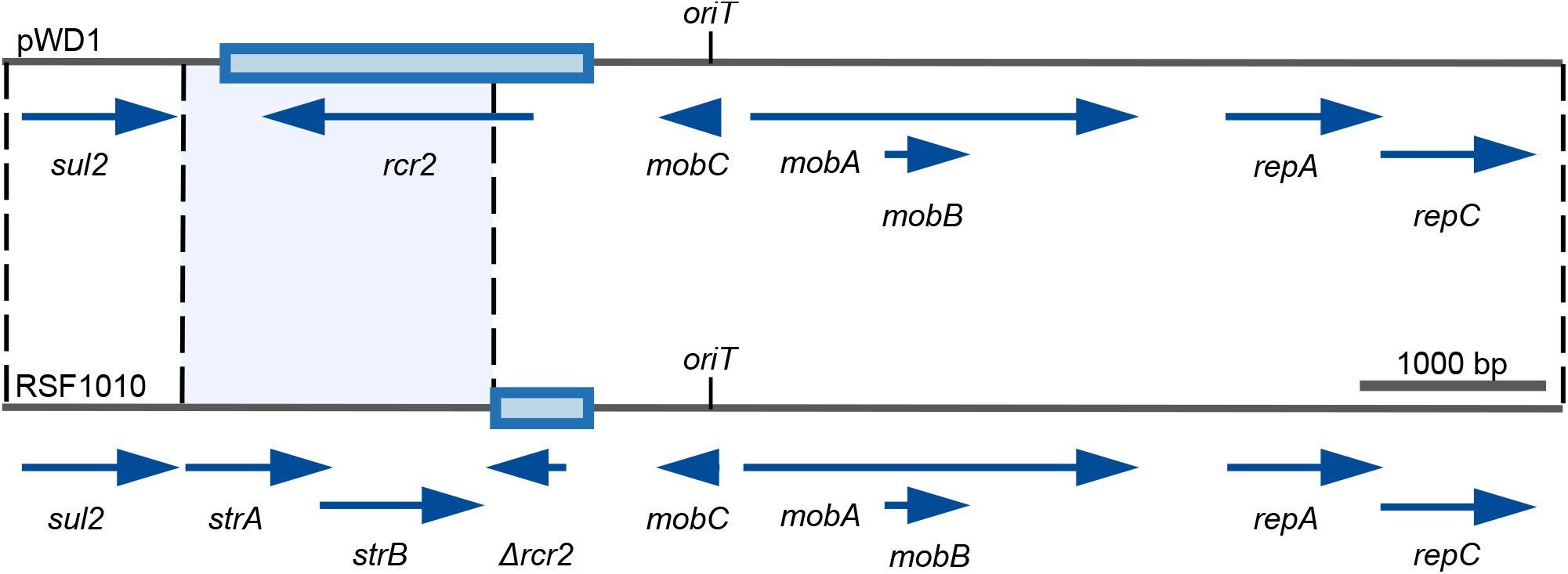
Comparison of plasmids pWD1 and RSF1010. Plasmid sequence is shown as a black line with the positions of genes indicated by labelled arrows below and the location of *oriT* shown above. The mobile element CR2 is shown as a thicker blue box. The light-blue shading highlights the region that differs between the plasmids and includes the *strAB* genes in RSF1010. Drawn to scale from GenBank accessions MW363525 and M28829 for pWD1 and RSF1010, respectively.

As metagenomic assemblies are often fragmented and plasmid replication genes may not be on the same contigs as ARGs that are carried on another region of the plasmid, we employed PlasFlow (22) to classify contigs in our metagenomic assembly as either chromosomal or plasmid. We identified a total of 93 plasmid contigs containing ARGs. The urban sediment samples contained significantly more plasmid contigs with ARGs than either of the rural sample types (Kruskal-Wallis; *P* < 0.001) whereas the urban water samples had significantly more ARG bearing plasmid contigs than the rural samples with no previous antibiotic use (Kruskal-Wallis*; P* < 0.05) (Figure 8). There was no significant difference in the number of ARG-containing plasmid contigs between rural samples with and without prior antibiotic use. Of the 93 contigs identified which contained ARGs, 78 contigs contained only one resistance gene with the remaining 15 contigs containing two or more ARGs (Table S3). All of the contigs that contained multiple resistance genes were found in urban samples and were closely related to known proteobacterial plasmids.

**Figure 8.**
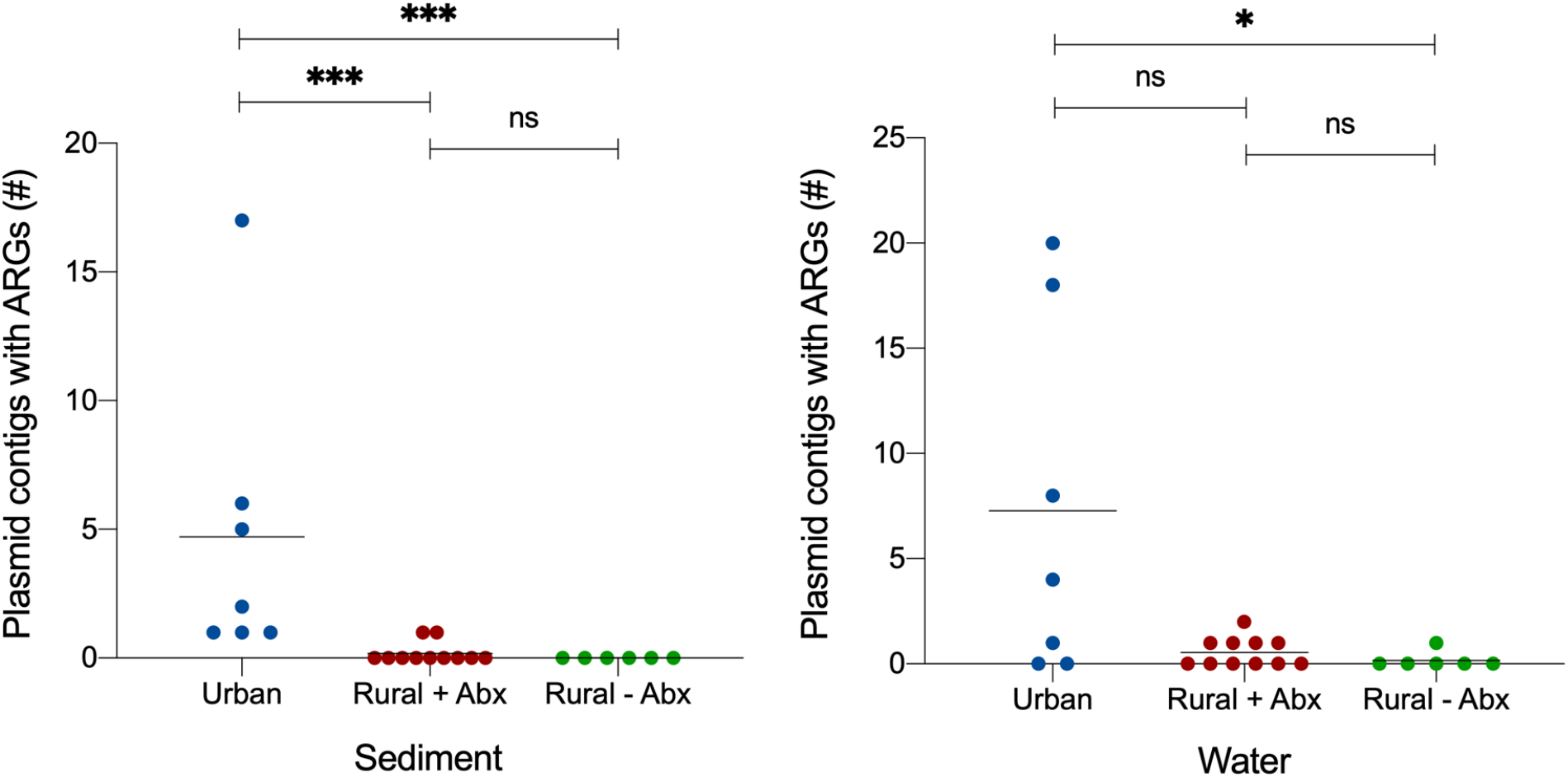
The number of contigs identified as plasmid by PlasFlow that carry an antibiotic resistance gene from Bangladesh surface water sites in sediment and rural, with and without antibiotic use (Abx), sediment and surface water samples. Kruskal-Wallis test: ^*^ *P* < 0.05 and ^***^ *P* < 0.001.

## Discussion

In this study we used quantitative culture and metagenomic techniques to understand the community composition and the level of antibiotic resistance genes in rural and urban surface water sites across Bangladesh. Selective plating showed that ESBL-producing coliforms were more prevalent in urban surface water compared to rural settings, consistent with reports of antibiotic resistant faecal coliforms in rivers across Asia (23, 24). However, the predictive value of the abundance of ESBL-producing coliforms for the total abundance of antibiotic resistance genes was found to be limited, suggesting that ESBL-producing coliforms are not necessarily a valid proxy to determine AMR load in environmental ecosystems.

In addition to quantitative culture of ESBL-producing coliforms, a metagenomic shotgun sequencing approach was used to characterise the microbiota of each sample and quantify the abundance of antibiotic resistance genes in water and sediment samples. We found that the water and sediment samples grouped together by their type (water or sediment) rather than the location they were collected from. Sediment samples were dominated by bacteria belonging to the genera *Pseudomonas*, *Azoarcus* and *Hydrogenophilacea* which is in line with other studies which have shown that sediment is dominated by the phylum Proteobacteria (25). Water samples were dominated by the cyanobacteria *Cyanobium* and *Microcystis* that cause harmful blooms in aquaculture ponds (26). *Microcystis* produces potent toxins which can kill fish but are also harmful to humans (27). The two river water samples and a public pond water sample collected in Dhaka clustered away from the other water samples and were defined by an increased abundance of bacteria associated with the human intestinal tract. The presence of increased amounts of the faecal indicator bacteria *E. coli* suggests that human waste is contaminating urban surface water (28).

Several different types of antibiotic were used in the rural aquaculture ponds which we surveyed (Table S1). The antibiotics were either mixed with feed or added directly to the ponds for the treatment of disease. Fluoroquinolone antibiotics such as ciprofloxacin and levofloxacin were the most widely used antibiotics in the rural aquaculture ponds, however high levels of fluoroquinolone resistance were not observed in the rural sites with prior antibiotic use. Resistance to fluoroquinolone drugs is mainly mediated by chromosomal mutations in the *parC* and *gyrA* genes, so the absence of dedicated resistance genes in these ecosystems may be unsurprising (29). However, we note that the multidrug efflux pump genes *mexV*, *mexF*, *adeI* and *adeH* were exclusively found in the rural sites with prior antibiotic use and these efflux systems are capable of exporting fluoroquinolones from the cell (30–33). In addition, other multidrug efflux pump genes capable of exporting fluoroquinolones such as *evgA* and *qacH* were found in these sites and in urban samples (34, 35). The macrolide drug erythromycin was another antibiotic which was widely used in aquaculture ponds that were sampled in this study. However, levels of macrolide resistance genes were low in the rural aquaculture ponds but extremely high in a subset of the urban samples. Notably, the erythromycin resistance gene *msrA* (36) was only present in the aquaculture ponds with prior antibiotic use. This gene was previously found in the intestinal contents of farmed rainbow trout and may thus be more commonly associated with aquaculture (37). Perhaps surprisingly, we did not observe a difference in the total load of antibiotic resistance genes in rural ponds with and without a history of antibiotic use. It may be possible that the widespread use of poultry manure as fish feed in both types of ponds (38–40) has introduced antibiotic resistant bacteria and/or antibiotics and could thus have minimised differences. Further research is needed to quantify the impact of these practices on the selection for antibiotic resistance in aquaculture ponds.

Our data suggest that antibiotic use in Bangladeshi aquaculture does not have a significant effect on the aggregated abundance of all antibiotic resistance genes in this ecosystem in comparison to urban surface water sites. Antibiotic resistance was the highest in urban areas which suggests that human factors contribute to the accumulation of antibiotic resistant bacteria in the environment. This was further corroborated by the correlation between the abundance of bacteria originating from the human gut and antibiotic resistance gene abundance observed in our study. The rivers and lakes of Dhaka are surrounded by areas with high population densities with 13.7% of households reporting that human waste is untreated and released directly into lakes, ponds or rivers (41). Our study thus extends on previous observations that link the introduction of human sewage into river and lake systems to high levels of antibiotic resistance genes (42).

By creating a metagenomic assembly of our short-read sequencing data we were able to identify contigs which contained plasmid replication initiation genes, contigs which could be circularised into complete plasmids and contigs which were predicted to be plasmids by PlasFlow and contained antibiotic resistance genes. We found that IncP, IncQ and various small plasmid types were most common. All of these plasmid types can replicate in a number of species belonging to the Enterobacteriaceae (43). Two complete plasmid sequences were recovered from the metagenomic assemblies of samples WCM1 and WD1. The small rolling-circle plasmid pWCM1 is closely related to plasmids such as pNMEC-O75D and p124_D that have been found in human and environmental *E. coli* isolates (44). The IncQ1 plasmid pWD1 is closely related and clearly ancestral to the well-characterised RSF1010. Although RSF1010 has been circulating globally since at least the 1960s, the structures of ancestral IncQ1 plasmids that only contain *sul2* have been predicted (45) but never found. The discovery of pWD1 in an urban water sample from 2018 is therefore surprising and demonstrates that this ancestral plasmid lineage has persisted stably for over 50 years. Due to the difficulties of assembling complete plasmid sequences from short-read metagenomic datasets, we were only able to circularise two plasmid sequences. For this reason, we also used additional methods to reconstruct plasmids revealing that urban samples had a higher number of plasmids carrying antibiotic resistance genes. This suggests that particularly in urban water bodies there exists an increased potential of horizontal gene transfer of mobile genetic elements carrying antibiotic resistance genes.

The microbiotas of surface water and sediment samples across Bangladesh are diverse, but antibiotic resistance genes are highly abundant in urban samples and are more commonly associated with plasmids in this setting. While the abundance of antibiotic resistance genes was considerably lower in rural than in urban settings, we nonetheless observed evidence for the selection for fluoroquinolone resistance mechanisms in ponds used for fish farming. Policies to minimise the use of antibiotics in aquaculture should thus remain a priority to reduce selection for antibiotic resistance. The presence of human gut bacteria was associated with high levels of antibiotic resistance genes, suggesting that contamination by human waste is an important driver for the presence of antibiotic resistance genes in surface water. Interventions aimed at improving access to clean water, sanitation and sewerage infrastructure may thus be important to reduce the risk of AMR dissemination in Bangladesh and other low- and middle-income countries.

## Materials and Methods

### Site selection

Paired surface water and sediment samples were collected in Bangladesh from 24 freshwater sites across three districts (Mymensingh, Shariatpur and Dhaka; Figure 1) in May and June of 2018. These sites spanned both rural and urban areas with different population densities. Samples were collected from 11 aquaculture ponds in the rural areas of two districts (Mymensingh and Shariatpur) with high commercial aquaculture activity. These ponds all had a history of antibiotic use within the past three months of collection. Six ponds with no history of antibiotic use were also sampled from these rural areas. In Mymensingh, 3 ponds used for domestic purposes were selected, while in Shariatpur, these were aquaculture ponds with no prior antibiotic use, which were used for culturing fingerlings. Antibiotic use information for the ponds was collected from local dealers who were responsible for supplying fish feed for these ponds. In addition to rural surface water sites, 7 water bodies (rivers, lakes and public ponds) were sampled in Dhaka. The public ponds were heavily used for domestic purposes and, while some had history of casual (non-commercial) fish cultivation, none of them had any prior antibiotic use.

### Sample collection

Samples were named using the following scheme; water (W) or sediment (S) followed by aquaculture (A) or control (C; ponds without antibiotic use). Sample sites were designated using (M) Mymensingh, (S) Shariatpur or (D) Dhaka and a number was included to differentiate samples. Further metadata on the samples, including temperature, pH and dissolved oxygen levels are provided in Table S1. Water samples were collected by submerging a sterile 500 ml Nalgene plastic bottle approximately 15 cm below the water’s surface. Bottles were capped before being removed from the water. The water samples were filtered through a 0.22 μm Sterivex-GP filter (Millipore) until water would no longer be passed through the filter. The filter units were then capped and stored in a cool box and transported to the laboratory within 12 hours of sampling. In addition to the water samples, approximately 10 g of sediment was taken from either the bed of the pond or from the bank 30 – 50 cm below the surface of the water. The sediment samples were stored in sterile 50 ml Falcon tubes and were transported with the water samples.

### Selective culturing for coliforms in surface water and sediment samples

Water and sediment samples were screened for the presence of Extended-Spectrum Beta-Lactamase (ESBL) producing coliforms by quantitative plating on Brilliance ESBL agar (Oxoid). Water and sediment samples were spread onto the plates and incubated for 48 hours at 37°C. In accordance with the manufacturer’s instructions blue, pink and green colonies were designated as coliforms and counted.

### DNA extraction and Illumina sequencing

DNA was extracted from the Sterivex filters and sediment samples using the DNeasy PowerWater Kit (Qiagen) and the DNeasy PowerSoil kit (Qiagen), respectively, in accordance with the manufacturer’s instructions. DNA concentrations were quantified using the Qubit dsDNA HS assay kit (Thermo Fisher) with all samples yielding more than 0.2 ng/μl. Negative control runs were performed for both kits by isolating DNA from sterile, distilled water: these yielded no detectable DNA. Metagenomic DNA libraries were prepared using the Nextera XT Library Prep Kit (Illumina). The libraries were pooled and sequenced on the HiSeq 2500 sequencing platform (Illumina) using a 150 bp paired-end protocol. Paired reads were adapter trimmed and both duplicates and reads less than 50 bp were removed using Trimmomatic 0.30 with Q15 as the sliding-window quality cut-off (46). The short-read sequencing data for this project has been deposited at the European Nucleotide Archive under accession number PRJEB39306.

### Taxonomic Profiling

To perform taxonomic profiling, the paired-end sequencing reads were mapped against clade specific markers using the MetaPhlAn2 package v.2.7.7 (47). The MetaPhlAn2 package was run with default parameters. The utility script merge_metaphlan_tables.py was used to merge all of the output files into a single tab delimited file.

### Source-sink analysis

Raw sequence reads from projects PRJNA254927, PRJEB7626 and PRJEB6092, which had previously been used as sources for source-sink analysis (48), were downloaded from the European Nucleotide Archive (ENA). These sequences represented freshwater, soil and gut metagenomes respectively. Adapters were removed from the sequence reads using fastp (49). Taxonomic counts were created for these metagenomic sequences and the 48 samples in this study by kraken2 v.2.0.9 (50) and Bracken v.2.6.0 (51) using a database containing bacterial, archaeal, viral and fungal sequences. A metadata table was created which described the environment that the sample was from and designated it as either a source or a sink. The taxonomic count table and the metadata table were used an input to the R package FEAST v.0.1.0 (19) which determined the proportion that each source contributed to each sink.

### Resistome profiling

Antibiotic resistance genes were identified using the ShortBRED package v.0.9.5 (52). The CARD database (53) (downloaded 1st July 2019) and the UniRef90 database (downloaded 4 July 2019) were used by ShortBRED-Identify to construct a marker database which the metagenomic reads could be mapped against. ShortBRED-Quantify.py was then used to map these paired-end reads against the database. The relative abundance in Reads Per Kilobase per Million reads (RPKM) was generated for each resistance gene family in the database. The RPKMs were summed for antibiotic resistance genes belonging to the same class and visualised with the pheatmap package (https://cran.r-project.org/web/packages/pheatmap/pheatmap.pdf) in R (54).

### Reconstruction of plasmids from metagenomic datasets

Metagenomic sequencing reads were assembled using the MEGAHIT v.1.1.3 assembler using default parameters (55). Contigs produced by MEGAHIT were then classified as plasmid or chromosomal by trained neural networks in the PlasFlow v1.1 program (22). Contigs designated to be of plasmid origin were queried against the CARD database by ABRicate v.0.9.8 (https://github.com/tseemann/abricate) to identify the presence of antibiotic resistance genes. Resistance genes were identified which had at least 95% identity and 50% coverage compared to the CARD database. Plasmid contigs were similarly queried against the PlasmidFinder database (20) to identify replication genes. Plasmids were circularised by comparing 300 bp from either end of putative plasmid-containing contigs using BLASTn (56). When ends were found to overlap, one copy of the overlapping sequence was removed to generate a complete, circularised plasmid sequence.

### Statistical analyses

The Shannon Diversity Index of the samples was calculated in R v.3.4.3 using the diversity function of the vegan package v.2.5-7 (57). Non-metric multidimensional scaling (NMDS) was also performed in R using the metaNMDS function of the vegan package. Permutational multivariate analysis of variance (PERMANOVA) was performed on a Bray-Curtis distance matrix of species abundance in R using the adonis function of the vegan package. Correlation between total ARG abundance and human gut bacterial contribution was calculated using the lm function in base R. Additional tests for determining statistical significance were performed as described in the text, implemented in GraphPad Prism v.8.3.1.

## Acknowledgments

This study was funded by the Royal Society through a Challenge Grant (CHG\R1\170015) and a Wolfson Research Merit Award (WM160092) to W.v.S. Metagenome sequencing was provided by MicrobesNG (http://www.microbesng.uk), which is supported by the BBSRC (grant number BB/L024209/1). R.S.M is funded by the Wellcome Trust Antimicrobials and Antimicrobial Resistance Doctoral Training Programme (215154/Z/18/Z).

## Author Contributions

W.V.S. and M.S.I. conceived this study. R.S.M., M.H.Z, I.T.A. and W.V.S. collected and processed the environmental samples with support from J.D.C. R.S.M., S.H. and R.A.M. analysed the data. R.S.M. and W.V.S. wrote the manuscript with input from all authors.

## Data Availability

Raw sequencing data have been submitted to the European Nucleotide Archive with accession number PRJEB39306.

