## Supplementary figures and images for "Microbiota analysis of rural and urban surface waters and sediments in Bangladesh identifies human waste as driver of antibiotic resistance"

### Figure S1

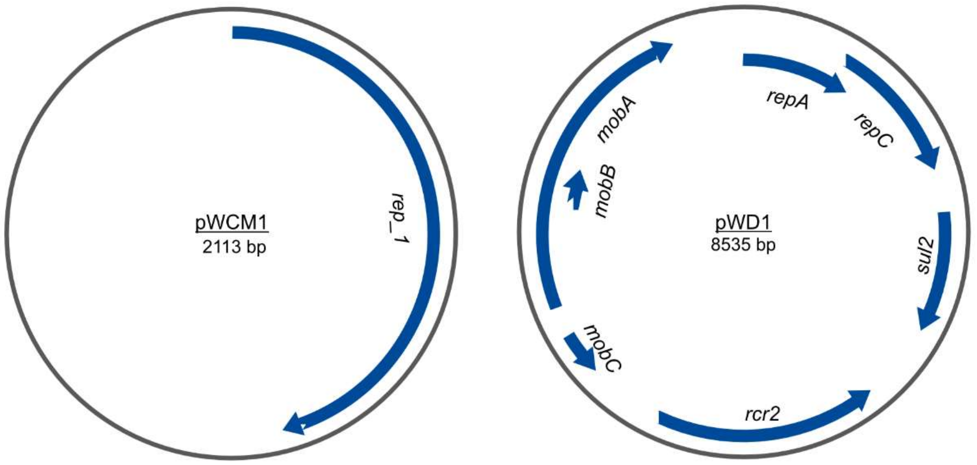


**Figure S1: Plasmid maps of pWCM1 (contig k141_206349) and pWD1 (k141_304072).**
