## Supplementary material for "Microbiota analysis of rural and urban surface waters and sediments in Bangladesh identifies human waste as driver of antibiotic resistance": Table S3

**Table S3. Contigs with one or more antibiotic resistance genes identified by PlasFlow as plasmids**

| Sample | Contig | Closest BLAST match in Genbank | Bacterial host of closest match | Identity (%) | Coverage (%) |
|---|---|---|---|---|---|
| **WCM3** | k141_292096 | pPm14C18 | *Proteus mirabilis* | 99.88 | 100.00 |
| **SD1** | k141_106889 | pSCU-397-2 | *Escherichia coli* | 99.11 | 99.00 |
| | k141_227167 | pAN70-1 | *Alcaligenes faecalis* | 99.96 | 100.00 |
| | k141_704017* | pNFYY023-1 | *Comamonas testosteroni* | 99.94 | 91.00 |
| | k141_836485 | pG5A4Y217 | *Escherichia coli* | 99.34 | 58.00 |
| | k141_99417 | pKP14812-MCR1 | *Klebsiella pneumoniae* | 97.57 | 40.00 |
| **WD1** | k141_128693 | p33 | *Escherichia coli* | 100.00 | 76.00 |
| | k141_134061 | Unnamed plasmid | *Butyricimonas faecalis* | 89.14 | 28.00 |
| | k141_139895 | pCP8-3-IncFIB | *Escherichia coli* | 95.43 | 99.00 |
| | k141_160447* | pCAV1335-92 | *Klebsiella oxytoca* | 98.76 | 100.00 |
| | k141_205613 | p1 | *Klebsiella pneumoniae* | 99.76 | 90.00 |
| | k141_256831* | pGENC284 | *Enterobacter hormaechei* subsp. *Xiangfangensis* | 100.00 | 100.00 |
| | k141_304072 | pSL7202-3 | *Salmonella enterica* subsp. *enterica* serovar *Typhimurium* | 99.98 | 81.00 |
| | k141_344452 | pRWC72a | Uncultured bacterium | 98.60 | 100.00 |
| | k141_377411* | pUCLAOXA232-5 | *Klebsiella pneumoniae* | 100.00 | 41.00 |
| | k141_391604 | pTZC1 | *Cutibacterium acnes* | 98.57 | 91.00 |
| | k141_44508 | pJF-786 | *Enterobacter cloacae* | 99.83 | 86.00 |
| | k141_467424* | pYH12207-3 | *Acinetobacter piscicola* | 100.00 | 92.00 |
| | k141_510896 | pG5A4Y217 | *Escherichia coli* | 99.46 | 51.00 |
| | k141_604491 | pSTN0717-64-1 | *Enterobacter hormaechei* | 99.75 | 100.00 |
| | k141_719363 | pSAN1-06-0624 | *Salmonella enterica* subsp. *enterica* serovar *Anatum* | 100.00 | 100.00 |
| | k141_719904 | pAeme6 | *Aeromonas media* | 99.77 | 100.00 |
| | k141_804869 | pMRGN207 | *Escherichia coli* | 99.59 | 70.00 |
| | k141_881861 | pBS228 | *Pseudomonas aeruginosa* | 99.94 | 92.00 |
| | k141_903595 | pWCX23_1 | *Aeromonas hydrophila* | 99.87 | 100.00 |

| | | | | | |
|---|---|---|---|---|---|
| | k141_91069 | pPN3F2_1 | *Shewanella aestuarii* | 99.23 | 68.00 |
| **SD2** | k141_324783 | pEC422_1 | *Escherichia coli* | 99.91 | 81.00 |
| | k141_325504 | Unnamed plasmid | *Klebsiella michiganensis* | 100.00 | 33.00 |
| | k141_343850 | RW109 | *Pseudomonas aeruginosa* | 99.92 | 58.00 |
| | k141_461478* | pAN70-1 | *Alcaligenes faecalis* | 100.00 | 100.00 |
| | k141_701410 | pGENC284 | *Enterobacter hormaechei* subsp. *Xiangfangensis* | 100.00 | 100.00 |
| | k141_743709 | pN1566_2 | *Salmonella enterica* subsp. *enterica* serovar *Schwarzengrund* | 99.77 | 100.00 |
| **WD2** | k141_109523 | pVB82_1 | *Acinetobacter baumannii* | 99.94 | 92.00 |
| | k141_113493* | pYH12207-3 | *Acinetobacter piscicola* | 99.93 | 78.00 |
| | k141_500987 | p24358-1 | *Salmonella enterica* subsp. *enterica* serovar *Bredeney* | 99.86 | 100.00 |
| | k141_723703* | pNFYY023-1 | *Comamonas testosteroni* | 100.00 | 99.00 |
| | k141_740211 | pVCGX2 | *Vibrio campbellii* | 99.94 | 100.00 |
| | k141_740911 | pMH17-012N_3 | *Citrobacter freundii* | 99.19 | 95.00 |
| | k141_76560 | p3 | Novosphingobium sp. *ES2-1* | 99.93 | 72.00 |
| | k141_777886 | p1681-tetX | *Empedobacter falsenii* | 98.40 | 98.00 |
| **SD3** | k141_1668271 | pOXA58_010030 | *Acinetobacter defluvii* | 99.82 | 100.00 |
| | k141_625842 | pCF39S | *Pseudomonas aeruginosa* | 99.94 | 99.00 |
| **SD4** | k141_546527 | pEI-2234-3 | *Edwardsiella ictaluri* | 61.00 | 100.00 |
| **SD5** | k141_806265 | pG5A4Y217 | *Escherichia coli* | 99.78 | 74.00 |
| **WD5** | k141_583117 | pGENC284 | *Enterobacter hormaechei* subsp. *Xiangfangensis* | 99.81 | 59.00 |
| **SD6** | k141_104572 | pAH01-4 | *Escherichia coli* | 99.74 | 100.00 |
| **WD6** | k141_124684 | p116753-FIIK | *Klebsiella pneumoniae* | 100.00 | 82.00 |
| | k141_158642* | pPm14C18 | *Proteus mirabilis* | 99.90 | 51.00 |
| | k141_173719 | pWP7-S18-ESBL-04 | *Klebsiella* sp. *WP7-S18-ESBL-04* | 99.97 | 72.00 |
| | k141_243506* | pYPR31 | *Providencia rettgeri* | 100.00 | 100.00 |
| **SD7** | k141_100623 | p63039 | *Myroides odoratimimus* | 99.89 | 100.00 |
| | k141_204766 | pKP20194a-p3 | *Klebsiella pneumoniae* | 100.00 | 100.00 |

| | | | | | |
|---|---|---|---|---|---|
| | k141_225349 | p1 | *Neisseria gonorrhoeae* | 98.74 | 100.00 |
| | k141_239404 | p24358-1 | *Salmonella enterica* subsp. *enterica* serovar *Bredeney* | 99.97 | 99.00 |
| | k141_24536 | pA2293-Ct2 | *Klebsiella pneumoniae* | 99.64 | 100.00 |
| | k141_270502 | pNA6 | Uncultured bacterium | 99.93 | 30.00 |
| | k141_333949 | pLraf_19_5_1 | *Lactococcus raffinolactis* | 94.90 | 37.00 |
| | k141_383255 | pHNCF11W-130kb | *Escherichia fergusonii* | 100.00 | 100.00 |
| | k141_393768 | pVCGX2 | *Vibrio campbellii* | 99.83 | 100.00 |
| | k141_451891 | pRErm46 | *Rhodococcus hoagii* | 99.00 | 89.00 |
| | k141_464290 | pC16KP0065-1 | *Klebsiella pneumoniae* | 100.00 | 39.00 |
| | k141_479837 | pEI-2234-3 | *Edwardsiella ictaluri* | 99.90 | 100.00 |
| | k141_556637 | pRGRH0399 | Uncultured bacterium | 94.84 | 80.00 |
| | k141_557329 | pHDC14-2.133K | *Enterococcus hirae* | 98.45 | 88.00 |
| | k141_569663* | pNFYY023-1 | *Comamonas testosteroni* | 99.98 | 80.00 |
| | k141_574437 | Plasmid 2 | *Salmonella enterica* subsp. *enterica* serovar *Typhi* | 99.44 | 96.00 |
| | k141_94024* | pAb-C63_1 | *Acinetobacter baumannii* | 99.95 | 79.00 |
| **WD7** | k141_113036 | pEI-2234-3 | *Edwardsiella ictaluri* | 100.00 | 100.00 |
| | k141_119966 | pHNCF11W-130kb | *Escherichia fergusonii* | 100.00 | 100.00 |
| | k141_139664 | pMS2H5VEB-1 | *Klebsiella pneumoniae* | 99.88 | 95.00 |
| | k141_16407 | pBS228 | *Pseudomonas aeruginosa* | 99.79 | 71.00 |
| | k141_174331 | pRSB222 | Uncultured bacterium | 97.35 | 83.00 |
| | k141_262055 | pG5A4Y217 | *Escherichia coli* | 90.98 | 85.00 |
| | k141_323094 | pN1566_2 | *Salmonella enterica* subsp. *enterica* serovar *Schwarzengrund* | 100.00 | 100.00 |
| | k141_337873 | pMH17-012N_3 | *CItrobacter freundii* | 98.83 | 92.00 |
| | k141_362572 | pALTS33 | Uncultured bacterium | 93.81 | 54.00 |
| | k141_38198 | pTet | *Campylobacter jejuni* subsp. *jejuni* | 99.89 | 100.00 |
| | k141_389765 | pCAP01 | *Capnocytophaga ochracea* | 89.11 | 35.00 |
| | k141_390832 | pMR0211 | *Providencia stuartii* | 99.96 | 95.00 |
| | k141_47250 | pE211-2 | *Enterococcus faecalis* | 99.41 | 100.00 |

| | | | | | |
|---|---|---|---|---|---|
| | k141_51571* | p3iANG | *Vibrio cholerae* | 97.91 | 85.00 |
| | k141_5646* | pYH12207-3 | *Acinetobacter piscicola* | 100.00 | 96.00 |
| | k141_7584 | p63039 | *Myroides odoratimimus* | 99.89 | 26.00 |
| | k141_93260* | pSY153-MDR | *Pseudomonas putida* | 99.97 | 96.00 |
| | k141_98708 | p345-185 | *Vibrio harveyi* | 94.32 | 77.00 |
| **SAS1** | k141_249579 | p3 | *Novosphingobium* sp. *ES2-1* | 99.92 | 85.00 |
| **WAS1** | k141_49220 | p4130-KPC | *Pseudomonas aeruginosa* | 99.93 | 100.00 |
| **SAS2** | k141_781158 | pEI-2234-3 | *Edwardsiella ictaluri* | 100.00 | 100.00 |
| **WAS3** | k141_1155247 | p3 | *Novosphingobium* sp. *ES2-1* | 90.90 | 92.00 |
| **WAM1** | k141_388101 | pP72_e | *Phaeobacter inhibens* | 98.95 | 99.00 |
| | k141_448754 | pEI-2234-3 | *Edwardsiella ictaluri* | 99.65 | 100.00 |
| **WAM3** | k141_457349 | pCF39S | *Pseudomonas aeruginosa* | 100.00 | 88.00 |
| **WAM6** | k141_67005 | pHH2-227 | Uncultured bacterium | 99.94 | 99.00 |
