## Supplementary material for "Microbiota analysis of rural and urban surface waters and sediments in Bangladesh identifies human waste as driver of antibiotic resistance": Table S2

**Table S2: Features of plasmid contigs.**

| Sample | Contig ID[a] | Rep type | Length (bp) | Coverage (%)[b] | Identity (%)[b] |
|---|---|---|---|---|---|
| **WCM1** | k141_156157 | Col156 | 5212 | 100 | 94.81 |
| | k141_206349* | Col(BS512) | 2254 | 100 | 100 |
| | k141_35625 | Col8282 | 3712 | 100 | 80.88 |
| **WD1** | k141_304072* | IncQ1 | 8676 | 100 | 100 |
| | k141_320207 | IncQ | 1420 | 51.33 | 88.31 |
| | k141_593572 | repUS43 | 1262 | 50.41 | 96.05 |
| | k141_711213 | IncQ1 | 24244 | 78.39 | 77.48 |
| | k141_728573 | Col(pWES) | 1941 | 93.26 | 80.95 |
| **WD2** | k141_452869 | IncQ | 2296 | 78.39 | 77.48 |
| **WD7** | k141_315908 | IncP6 | 1166 | 100 | 99.63 |
| | k141_77466 | IncQ | 4072 | 78.39 | 77.48 |

[a] Plasmid contig which could be circularised.

[b] Coverage and identity are of the closest *rep* gene in the PlasmidFinder database.
