## Supplementary material for "Microbiota analysis of rural and urban surface waters and sediments in Bangladesh identifies human waste as driver of antibiotic resistance": Table S1

**Table S1: Metadata of surface water samples**

| Sample[a] | pH | Dissolved Oxygen (%) | T (°C) | Surface water type | Fish[b] | Antibiotics used[c] | Date collected |
|---|---|---|---|---|---|---|---|
| **WAM1** | 7.8 | 76.8 | 26.4 | Commercial aquaculture | Shing (fingerlings) | C | 08/05/2018 |
| **WAM2** | 7.7 | 89.7 | 27.9 | Commercial aquaculture | Shing | E, S, T | 08/05/2018 |
| **WAM3** | 7.6 | 73.1 | 27.8 | Commercial aquaculture | Koi | C, O | 08/05/2018 |
| **WAM4** | 7.6 | 37.9 | 27.1 | Commercial aquaculture | Pabda | C, E, S, T | 08/05/2018 |
| **WAM5** | 7.3 | 73.8 | 27.3 | Commercial aquaculture | Pangas | C, S | 08/05/2018 |
| **WAM6** | 7.1 | 59.9 | 28.0 | Commercial aquaculture | Pangas, Shing | C, E, S, T | 08/05/2018 |
| **WCM1** | 7.7 | 70.6 | 26.5 | Household pond | NA | NA | 08/05/2018 |
| **WCM2** | 8 | 97.1 | 26.8 | Household pond | NA | NA | 08/05/2018 |
| **WCM3** | 9.2 | 126.8 | 27.0 | Household pond | NA | NA | 08/05/2018 |
| **WAS1** | 7.5 | 68.3 | 32.1 | Commercial aquaculture | Rui, Catla, Mrigal | E | 11/06/2018 |
| **WAS2** | 8.4 | 112 | 32.1 | Commercial aquaculture | Rui, Catla, Mrigal | C, E, S | 11/06/2018 |
| **WAS3** | 8.3 | 102.6 | 34.2 | Commercial aquaculture | Rui, Catla, Mrigal | C | 11/06/2018 |
| **WAS4** | 8.9 | 135.2 | 35.9 | Commercial aquaculture | Rui, Catla, Mrigal | E, S | 11/06/2018 |
| **WAS5** | 7.8 | 66.8 | 35.7 | Commercial aquaculture | Rui, Catla, Mrigal | C, S, T | 11/06/2018 |
| **WCS1** | 8.1 | 85.2 | 31.6 | Commercial aquaculture (no antibiotic use) | Rui, Catla, Mrigal (fingerlings) | NA | 11/06/2018 |
| **WCS2** | 7.4 | 44 | 31.3 | Commercial aquaculture (no antibiotic use) | NA | NA | 11/06/2018 |
| **WCS3** | 7.5 | 69.3 | 34.1 | Commercial aquaculture (no antibiotic use) | NA | NA | 11/06/2018 |
| **WD1** | 7.1 | 6 | 26.8 | Urban lake | NA | NA | 11/05/2018 |
| **WD2** | 7.4 | 15.4 | 28.0 | Urban lake | NA | NA | 11/05/2018 |
| **WD3** | 7.6 | 67.7 | 32.5 | Urban pond | NA | NA | 11/05/2018 |
| **WD4** | 7.4 | 78.4 | 32 | Urban lake | NA | NA | 11/05/2018 |
| **WD5** | 7.8 | 76 | 30.5 | Urban pond | NA | NA | 11/05/2018 |
| **WD6** | 7.1 | 12.3 | 28.2 | Urban river | NA | NA | 18/05/2018 |
| **WD7** | 7.1 | 9.5 | 28.3 | Urban river | NA | NA | 19/05/2018 |

*a*

[a] Surface water samples are coded as follows. W identifies them as water samples, the second letter identifies whether the samples were collected in rural settings (A: aquaculture, C: control, non-aquaculture) or urban sites (D: Dhaka). In the samples collected at rural sites, the region is indicated with the final letter (M: Mymensingh, S: Shariatpur).

[b] Binomial names: Shing: *Heteropneustes fossilis*, Koi: *Anabas cobojius,* Pabda: *Callichrus pabda*, Pangas: *Pangasius pangasius*, Rui*: Labeo rohita*, Catla: *Catla catla*, Mrigal: *Cirrhinus cirrhosis*; NA: not applicable

[c] C: ciprofloxacin, E: erythromycin, S: sulfadiazine, T: trimethoprim, O: oxytetracycline; NA: not applicable.
